# Arterial carboxyhaemoglobin levels in children admitted to PICU: a retrospective observational study

**DOI:** 10.1101/490334

**Authors:** Ankur Chawla, Samiran Ray, Adela Matettore, Mark J Peters

## Abstract

While carbon monoxide (CO) is considered toxic, low levels of endogenously produced CO are protective against cellular injury induced by oxidative stress. Carboxyhaemoglobin (COHb) levels have been associated with outcomes in critically ill adults. We aimed to describe the distribution of carboxyhaemoglobin in critically ill children and the relationship of these levels with clinical outcomes. This retrospective observational study was conducted at a large tertiary paediatric intensive care unit (PICU). We included all children admitted to the PICU over a two-year period who underwent arterial blood gas analysis. We measured the following: Population and age-related differences in COHb distribution; (ii) Change in COHb over the first week of admission using a multi-level linear regression analysis; (ii) Uni- and multivariable relationships between COHb and length of ventilation and PICU survival. Arterial COHb levels were available for 559/2029 admissions. The median COHb level was 1.20% (IQR 1.00-1.60%). Younger children had significantly higher COHb levels (p-value <2 x 10^−16^). Maximum Carboxyhaemoglobin was associated with survival 1.67 (95% CI 1.01-2.57, p-value=0.02) and length of ventilation (odds ratio 5.20, 95% CI 3.07-7.30, p-value 1.8 x 10^−6^) following multi-variable analysis. First measured and minimum COHb values were weakly associated with length of ventilation, but not survival. In conclusion, children have increased COHb levels in critical illness, which are greater in younger children. Higher COHb levels are associated with longer length of ventilation and death in PICU. This is likely to reflect increased oxidative stress.

## Introduction

Exogenous carbon monoxide (CO) has long been considered toxic due to competitive inhibition of haemoglobin oxygen binding. However, endogenous CO is increasingly being associated with beneficial physiological effects [1]. Heme is oxidised to produce CO, free iron and biliverdin, a reaction catalysed by heme oxygenase (HO). Of the three known isoforms of HO, HO-1 is highly induced by oxidative stress, it acts to limit damage from the oxidant heme [2]. CO therefore may be a biomarker of oxidative stress. Carbon monoxide can activate soluble guanylate cyclase, and hence the cyclic guanylate monophosphate (cGMP) signalling pathway. This is thought to be a common signalling pathway for CO’s anti-apoptotic, anti-inflammatory and vaso-active properties [3,4]. Protective effects of CO have been demonstrated in laboratory models of sepsis, acute lung injury, organ transplantation and reperfusion injury [5–8]. In vivo, CO is mostly eliminated in exhaled air, but a proportion remains bound to haemoglobin – this is determined by the rate of CO production and minute ventilation [9].

Elevated levels of exhaled CO are associated with smoking, and patients with chronic respiratory disease [10]. This suggests that exhaled CO may be a marker of lung injury. Elevated exhaled CO levels have also been described in critically ill adults in the intensive care unit (ICU) [11]. However, exhaled CO levels were correlated with arterial carboxyhaemoglobin (COHb) and serum bilirubin levels, suggesting increased CO production due to increased in systemic HO activity rather than lung injury in critical illness [12].

Arterial COHb levels have previously been associated with outcomes in adult ICU patients. Melley et al described a U-shaped relationship between ICU survival and COHb levels: risk adjusted mortality was associated with a lower minimum and greater maximum COHb level in patients post cardio-pulmonary bypass [13]. The authors hypothesised that an optimal amount of CO conferred anti-inflammatory effects in critically ill patients: patients unable to mount a HO-1 response to stress (e.g. due to genetic polymorphisms) may lack this anti-inflammatory response. In patients with high COHb levels, high levels of oxidative stress may account for the increased mortality. Alternatively, the deleterious effects of concomitant high levels of free iron, itself reactive, may account for negative outcomes [14].

Fazekas et al similarly demonstrated the association between low COHb and mortality in adult medical ICU patients [15]. A small study of COHb levels in preterm neonates did not find associations between the presence and absence of sepsis but did find associations between postnatal age and haemoglobin levels [16]. However, this is likely to be related to high levels of haemolysis in the early neonatal period. There are no studies of COHb levels in critically ill children. Increased HO-1 activity in children has been described in disease states such as obesity [17] and haemolysis [18]. Therefore, similar responses as seen in adult ICU patients should be expected.

In this retrospective, observational study, we aimed to 1) evaluate the distribution of arterial COHb in children admitted to a medical paediatric intensive care unit; 2) describe the trajectory of arterial COHb levels over the first 7 days of admission; 3) seek associations between arterial COHb and ICU outcomes: given the low rate of mortality on paediatric ICU, we also sought associations between COHb and length of ventilated stay.

## Methods

We conducted a retrospective, observational study using data from a single centre, 23-bedded general (non-cardiac) paediatric intensive care unit (PICU). Data were collected from the electronic health records (Philips Intellivue Critical Care and Anaesthesia, Philips Electronics, Amsterdam, the Netherlands) for all children following unplanned admission to PICU who underwent arterial blood gas analysis between 1^st^ January 2015 and 31^st^ December 2016. All children from neonates to 16 years were included. Arterial blood gas analysis was carried out using the ABL 90 Flex Blood Gas analyser (Radiometer Medical, Bronshoj, Denmark). COHb is expressed as a fraction of total haemoglobin concentration, measured by co-oximetry. As a majority of children undergo capillary sampling, with a potential of haemolysis at the point of sampling, only arterial gas samples were considered. Venous samples were also excluded - although arterio-venous correlation for COHb is excellent, the mean difference arterio-venous of 0.15% (95% CI 0.13-0.45%) [19] may have confounded results for children with low COHb levels. Children who underwent continuous veno-venous haemofiltration (CVVH) were also excluded due to the likelihood of extra-corporeal haemolysis.

Time of collection and COHb level for each arterial gas sample analysed were collected. For each patient who underwent arterial blood gas analysis, admission and discharge date and time, minute ventilation, length of ventilation in days, status at discharge, age in months, sex and Paediatric Index of Mortality-3 (PIM-3) score data were collected.

Continuous data were described as median and inter-quartile intervals. Differences between age groups (<1 month, 1 month-1 year and >1 year of age) were analysed using the Kruskal-Wallis test. The mean change in COHb over the first week of admission was evaluated following multi-level regression analysis, using COHb as the dependent variable, day of admission, age, PIM-3 score and the minute ventilation as the fixed effect variables and the patient and admission identifier as the random effect variables. The first COHb level following PICU admission, maximum and minimum COHb levels during admission were analysed for relationship with length of ventilation and survival status at discharge. Univariate associations with COHb levels were sought using Spearman’s correlation for length of ventilation, and Mann-Whitney tests for survival at discharge. The predictive value for survival was also quantified using receiver operating characteristic (ROC) curves. Linear and logistic regression analysis for length of ventilation and survival status at discharge were used respectively. Age and PIM-3 score were used as confounders, with length of ventilation added as a confounder for survival status. All data were analysed using Microsoft Excel (Microsoft Corp, Redmond, WA) and r (www.r-cran.org).

The study was reviewed and approved by the institutional research department (ref 17BB36). As the investigators only accessed previously collected, non-identifiable information/data under GAfREC 2011, the need for individual patient consent was waived by the institutional research governance department.

## Results

Over the 2-year period, there were 2027 admissions to PICU, 1239 (61.1%) of which were unplanned. Of these, 619 (50.0%) had arterial blood gas analysis during their admission. Sixty-one children required CVVH, leaving 558 (45.0%) children for further analysis. The baseline characteristics of these children are shown in table 1.

**Table 1:**
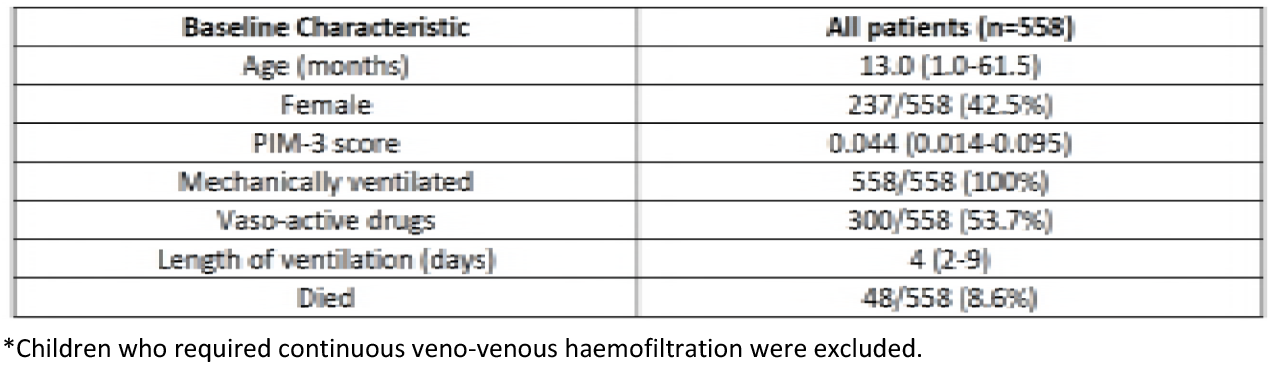
Baseline characteristics of children following unplanned admissions to PICU who underwent arterial blood gas analysis in 2015-16. Continuous variables expressed as medians and interquartile ranges and categorical variables expressed as fractions and percentages.

| Baseline Characteristic | All patients (n=558) |
| --- | --- |
| Age (months) | 13.0 (1.0-61.5) |
| Female | 237/558 (42.5%) |
| PIM-3 score | 0.044 (0.014-0.095) |
| Mechanically ventilated | 558/558 (100%) |
| Vaso-active drugs | 300/558 (53.7%) |
| Length of ventilation (days) | 4 (2-9) |
| Died | 48/558 (8.6%) |

\*Children who required continuous veno-venous haemofiltration were excluded.

### Distribution and age-related differences in COHb

The distribution of COHb values over the length of the admission is shown in Figure 1. The median value was 1.20% (IQR 1.00-1.60%). The median of the first value measured on PICU was 1.00% (IQR 0.70-1.30%) and of the maximum value during admission was 1.40% (IQR 1.20-2.00%); while of minimum values it was 0.80% (IQR 0.60-1.10%).

**Fig 1:**
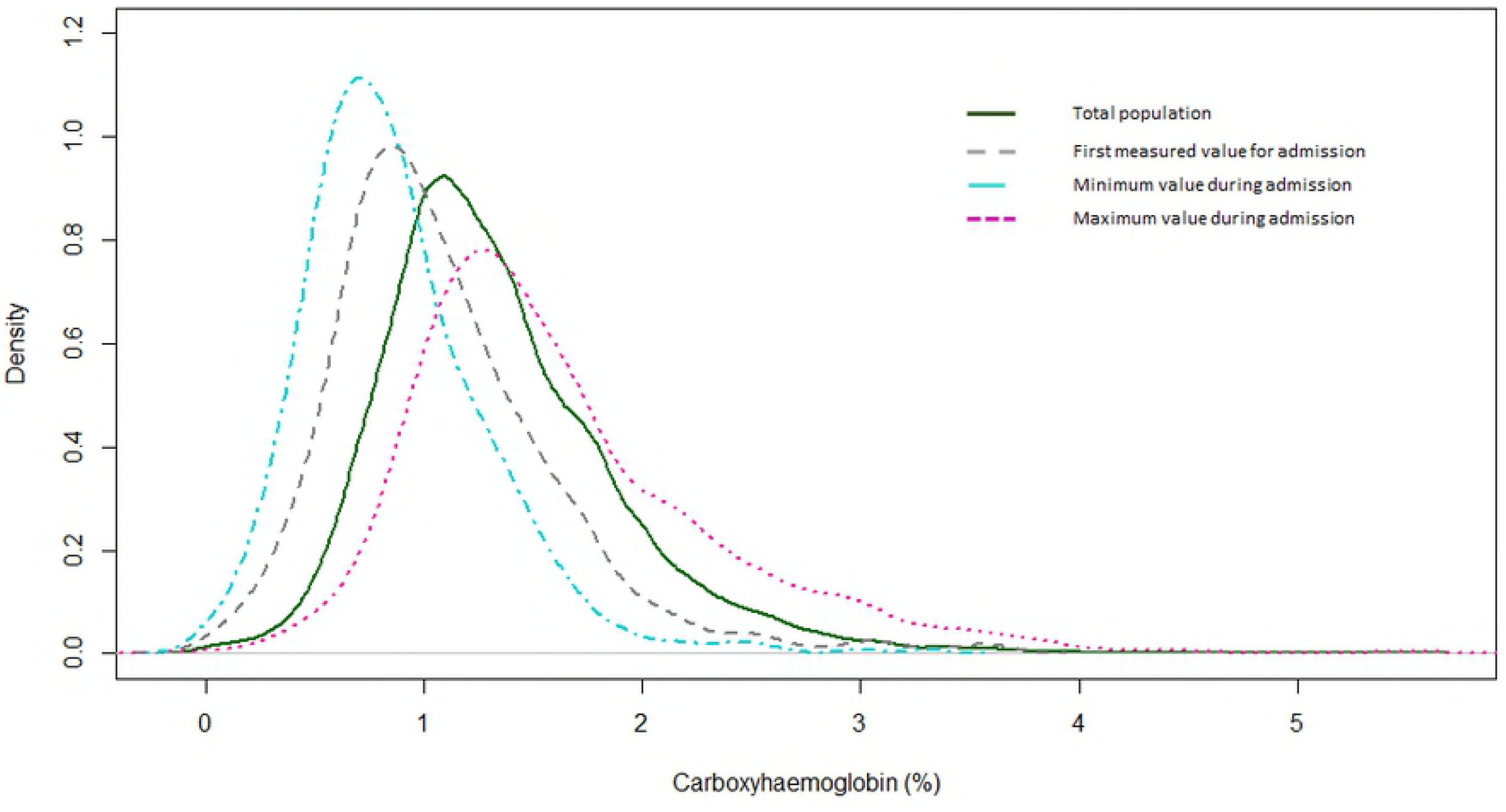
Distribution of carboxyhaemoglobin (COHb) for children admitted to the paediatric intensive care unit in 2015-16 who underwent arterial blood gas analysis. The continuous line shows the overall distribution for the cohort. The grey dashed line shows the distribution for the first measured COHb value; the pink dotted line shows the distribution of maximum values during admission and the blue dashed line the minimum values during admission.

Age was categorised into children less than 1 month, between 1 month and 1 year, 1 to 5 years and 5-16 years, reflecting the proportion of admissions both locally and nationally. The COHb values were significantly different between age categories (Kruskall-Wallis chi statistic 538.8, p-value <2 x 10^−16^). Younger children had wider variation in COHb levels with a tendency towards higher values (Figure 2a). To ensure this was not explained by once category having arterial blood samples for a longer period of time, we also compared first COHb values. These also significantly varied with age (Kruskal-Wallis chi statistic 47.1, p-value 3.2 x 10^−10^), with infants having higher values, and showing greater variation (Figure 2b).

**Fig 2:**
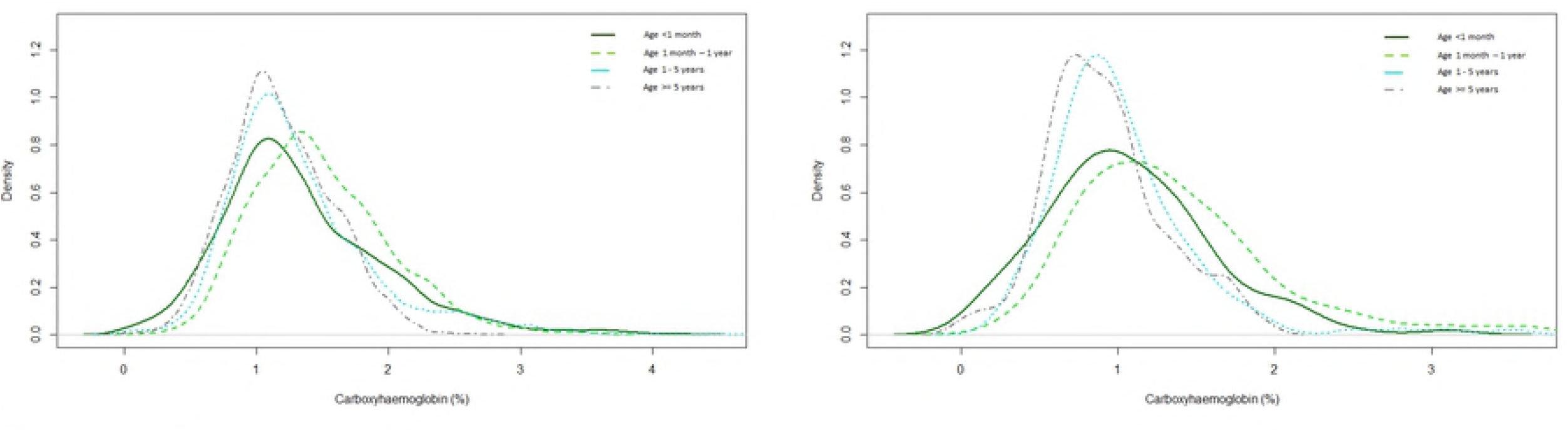
The distribution of (a) all carboxyhaemoglobin (COHb) values throughout PICU stay (left), and (b) the first measured value of COHb (right), according to age groups. There is a statistically significant difference between the groups for all COHb values (Kruskal-Wallis chi-squared 538.8, p-value <2 x 10^−16^) and first measured COHb values (Kruskal-Wallis chi-squared 47.1, p-value 3.2 x 10^−10^).

### Trajectory of COHb over the first 7 days of admission

The trajectory of COHb was evaluated using a multi-level regression model to control for each patient admission. COHb increased over the first week. The mean COHb level, controlling for PIM-3, age and minute volume, was 1.17 (95% CI 1.12-1.22) on day 0, compared to 1.43 (95% CI 1.38-1.48) on day 3 and 1.62 (95% CI 1.56-1.671) on day 7 (Table 2; Figure 3).

**Table 2:** Mean COHb levels for each day of admission in the first week following multi-level linear regression analysis. The regression model was constructed with COHb as the dependent variable, days, minute volume, PIM-3 score and age as the fixed effect variables and unique identifier for patient and admission as random effects variables. Day 0 is defined as the first day of admission till midnight; following days are defined by the 24-hour period from midnight to 23:59.

| Day of admission | Number of measurements | Mean COHb (95% CI)<br>following multi-level regression<br>analysis |
| --- | --- | --- |
| Day 0 | 3272 | 1.17 (1.12-1.22) |
| Day 1 | 2199 | 1.27 (1.23-1.33) |
| Day 2 | 1738 | 1.38 (1.33-1.43) |
| Day 3 | 1243 | 1.43 (1.38-1.48) |
| Day 4 | 919 | 1.50 (1.45-1.55) |
| Day 5 | 662 | 1.52 (1.46-1.57) |
| Day 6 | 531 | 1.51 (1.45-1.56) |
| Day 7 | 428 | 1.62 (1.56-1.671) |

**Fig 3:**
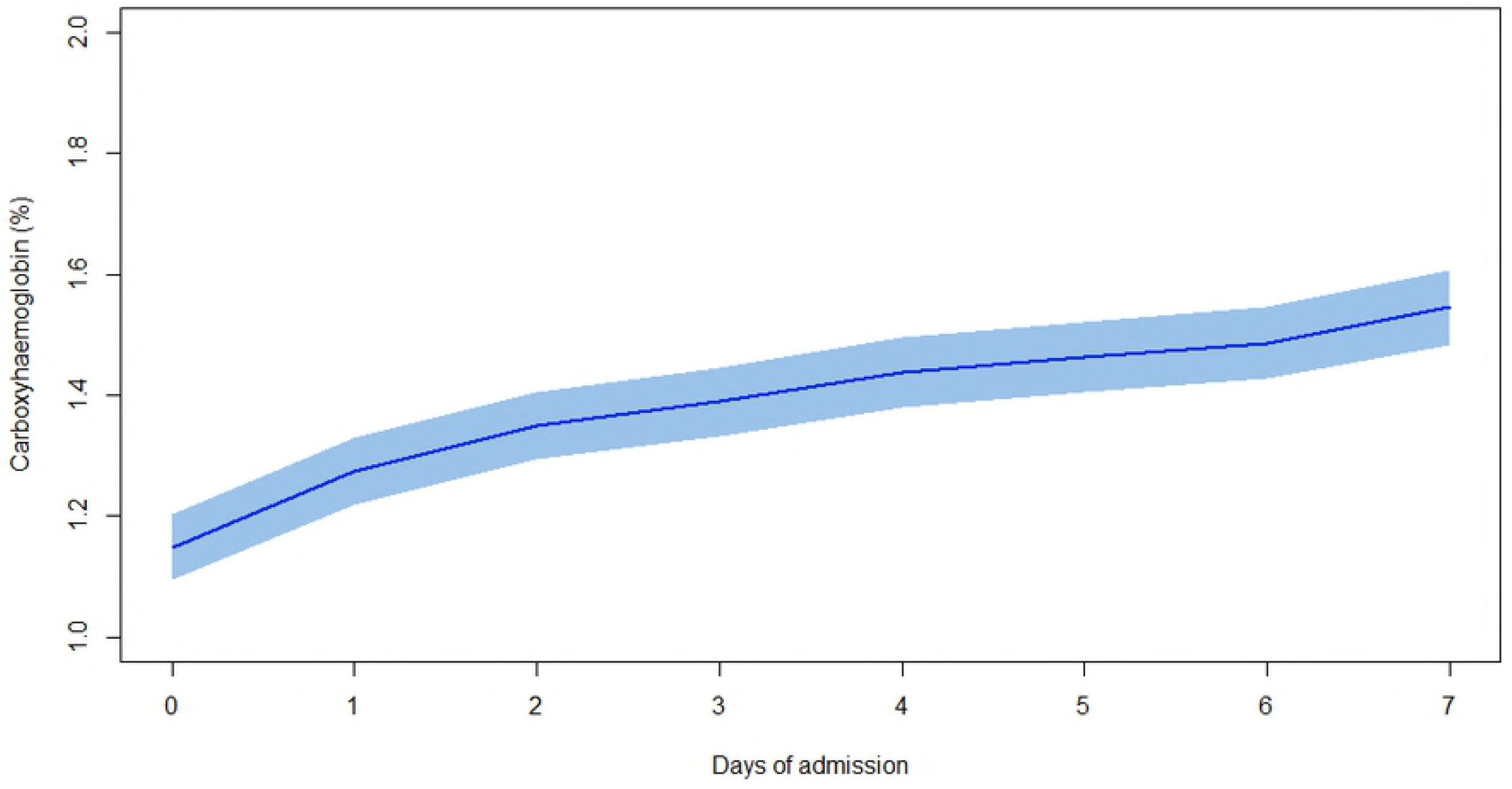
Mean COHb levels with 95% confidence intervals according to day of admission, following multi-level linear regression analysis. The COHb level rises over the first 7 days of admission. The regression model was constructed with COHb as the dependent variable, days, PIM-3 score, age and minute ventilation as the fixed effect variables and unique identifier for patient and admission as random effects variables. Day 0 is defined as the first day of admission till midnight; following days are defined by the 24-hour period from midnight to 23:59.

### Association between COHb and PICU outcomes

Associations between COHb and PICU outcomes were sought for (i) first measured COHb values, (ii) maximum COHb values during admission, and (iii) minimum COHb values.

i. The first and minimum COHb values were not significantly different between children who survived and those who died at PICU discharge (Mann-Whitney p-value 0.70 for first recorded COHb; 0.36 for minimum COHb). The maximum COHb was significantly higher in children who died (median 1.60, IQR 1.30-1.182) compared to those who survived (median 1.40, IQR 1.10-1.90; Mann-Whitney p-value 0.04). (Figure 4) The first measured COHb value poorly predicted survival at ICU discharge, with an area under the ROC curve of 0.52. Following logistic regression, with age, PIM-3 score and length of ventilation as confounders, first COHb did not independently predict survival at discharge. The first measured COHb value was poorly correlated with the length of ventilation (Figure 5), with a Spearman’s correlation coefficient of 0.28. Following linear regression with age and PIM-3 as confounders, first COHb was significantly associated with length of ventilation, with an odds ratio of 5.2 (95%CI 2.4-8.0, p-value 0.0003). However only 2.3% of the variation in length of ventilation was accounted for by the three variables (i.e. R^2^ for COHb, age and PIM-3).
ii. The maximum COHb value predicted survival status slightly better, with maximum values being higher in the non-survivor group (Figure 4). The area under the ROC curve was only 0.59, although after controlling for PIM-3 and age, the odds ratio for maximum COHb was 1.67 (95% CI 1.01-2.57, p-value=0.02). The maximum COHb was poorly correlated with length of ventilation (Spearman R 0.41) (Figure 5). Following multivariable analysis, maximum COHb was independently associated with length of ventilation, with an odds ratio of 5.20 (95% CI 3.07-7.30, p-value 1.8 x 10^−6^). The model R^2^ was 0.04, only slightly better than that for first COHb
iii. The minimum COHb value poorly predicted survival status (Figure 4) with a correlation coefficient of 0.14. Minimum COHb did not predict survival after multivariable analysis. Minimum COHb poorly predicted length of ventilation (Figure 5), both on uni-variable (Spearman R 0.03) and multi-variable analysis.

**Fig 4:**
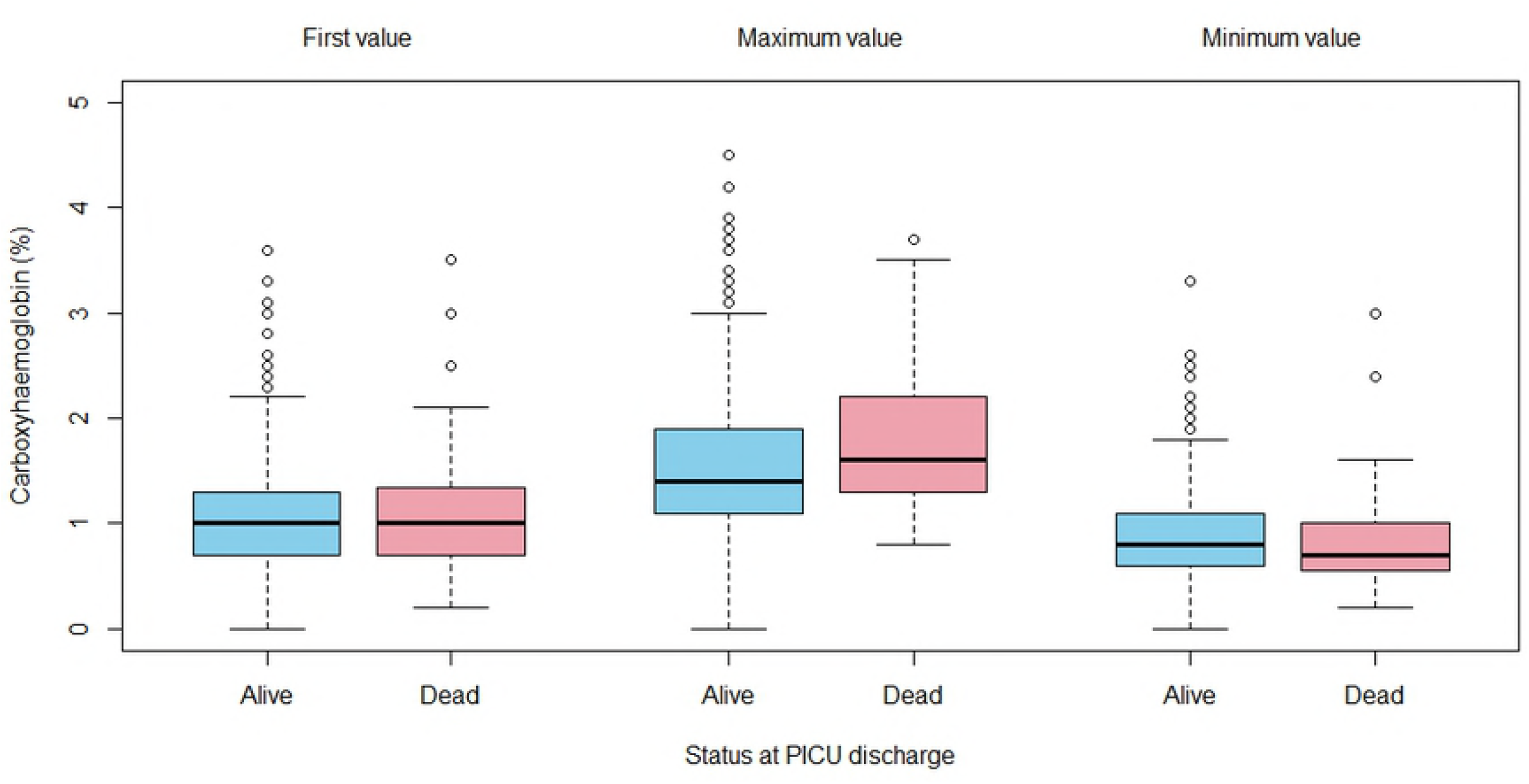
Box plot of measured COHb values and survival status at PICU discharge. The first, minimum and maximum COHb values (%) and the survival status at PICU discharge are represented. Neither the first nor the minimum COHb values predicted survival status at ICU discharge. The maximum COHb values were slightly better at predicting survival status at PICU discharge.

**Fig 5:**
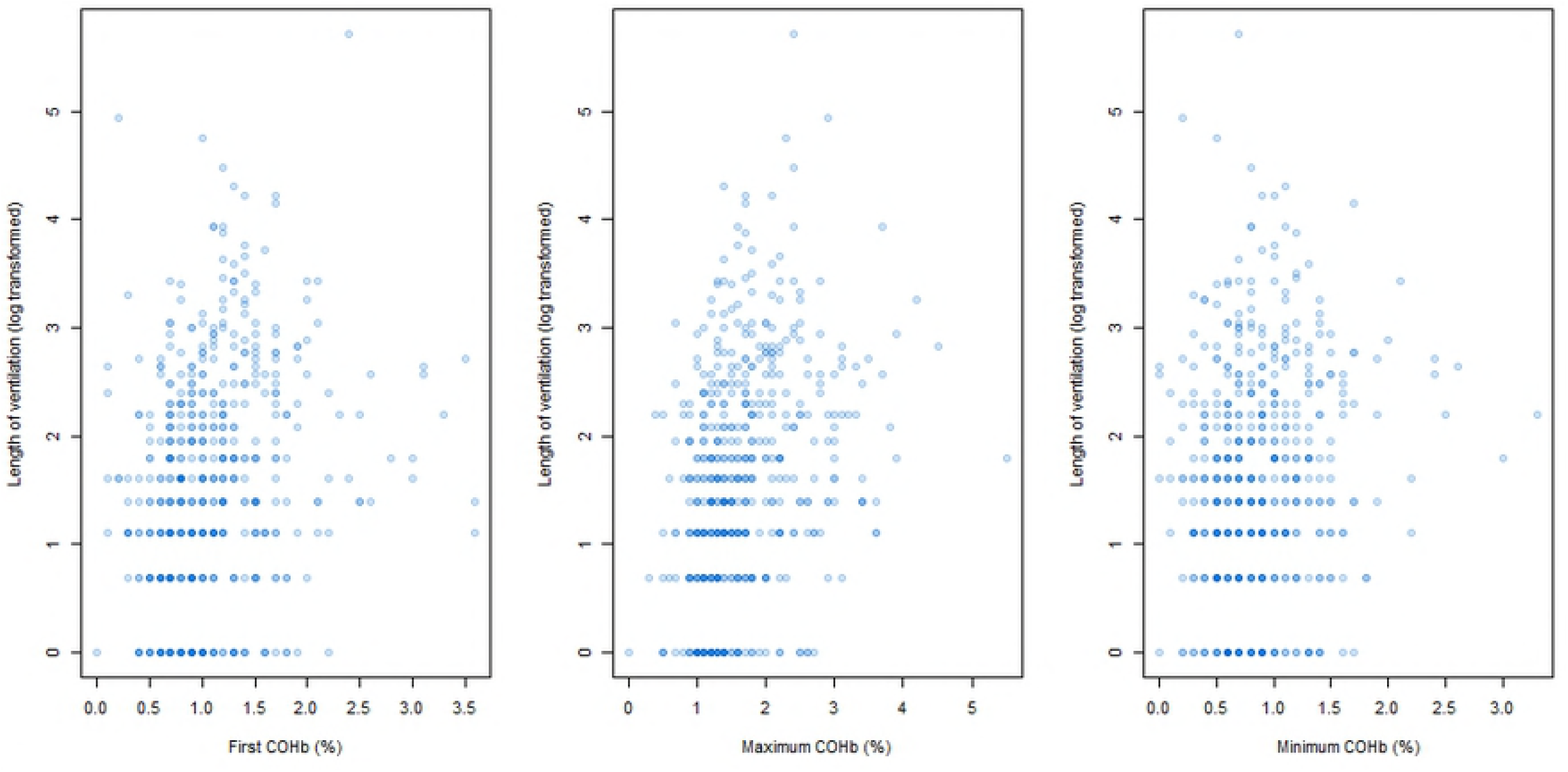
Scatter plot of measured COHb values and length of ventilation (log transformed). The first, minimum and maximum COHb values (%) and the length of ventilation in hours are represented.

## Discussion

In this single centre retrospective observational study, we describe the distribution of arterial COHb in children admitted to ICU. COHb levels rise with time mechanically ventilated. There is poor association between mortality and maximum COHb during PICU admission, and better association with length of ventilation. The association with length of ventilation extends to the first measured COHb and the minimum COHb, both positively correlated, albeit weakly.

The distribution of COHb was similar to that previously described in adult critical illness (Melley and Fazekas described time-weighted mean values of 1.2 and 1.6% respectively, similar value in our cohort is 1.3%). Younger children in our cohort demonstrated higher and more variable COHb levels. This may reflect severity of illness: there was a significant difference in the distribution of PIM scores, (median PIM scores: <1 month 0.078; 1 month- 1 year 0.044; 1 year-5 years 0.037; >5 years 0.035, Kruskal-Wallis chi-squared 42.7, p-value 2.9 x 10^−9^), although the COHb values were highest in the 1 month to 1 year cohort. It is possible that neonates are less able to mount the same degree of HO-1 response according to disease severity, this needs further study.

Our data also demonstrated that COHb increases with time. It is therefore unsurprising that the maximum COHb correlates with length of ventilation and mortality: the maximum value is likeliest to occur prior to death. In our cohort this is confounded by the use of arterial samples only: unlike in adult ICU, arterial catheters are likely to be in place only if the child is still ventilated and critically unwell. It is quite possible that in children who are improving and are extubated, the COHb falls. The rise in COHb may be in response to ongoing inflammation from ventilator associated lung injury, or hyperoxia: HO-1 induction has been described in response to hyperoxic damage [5]. We have previously described a bias towards higher saturation targets in our centre [20], raising this mechanistic possibility.

Melley et al have previously described a U-shaped relationship between COHb and mortality in the ICU in a cohort with low mortality. This was not replicated in a general adult ICU population with higher mortality. Although mortality was weakly associated with maximum COHb in our cohort, there was no association with minimum COHb. However, as COHb rises over time, and the sickest children in PICU die early [21,22], it may be that any U-shaped relationship is subtler. In order to test this further post-hoc, the risk adjusted mortality according 0.5% intervals of maximum COHb values were calculated, relative to the risk adjusted mortality of the median category (i.e. 1.0-1.5%). Lower maximum COHb values however had lower risk adjusted mortality (Figure 6), refuting such a U-shaped relationship in this cohort. However, this may be confounded by the size of the cohort, and the limitation to using only arterial COHb values.

**Fig 6:**
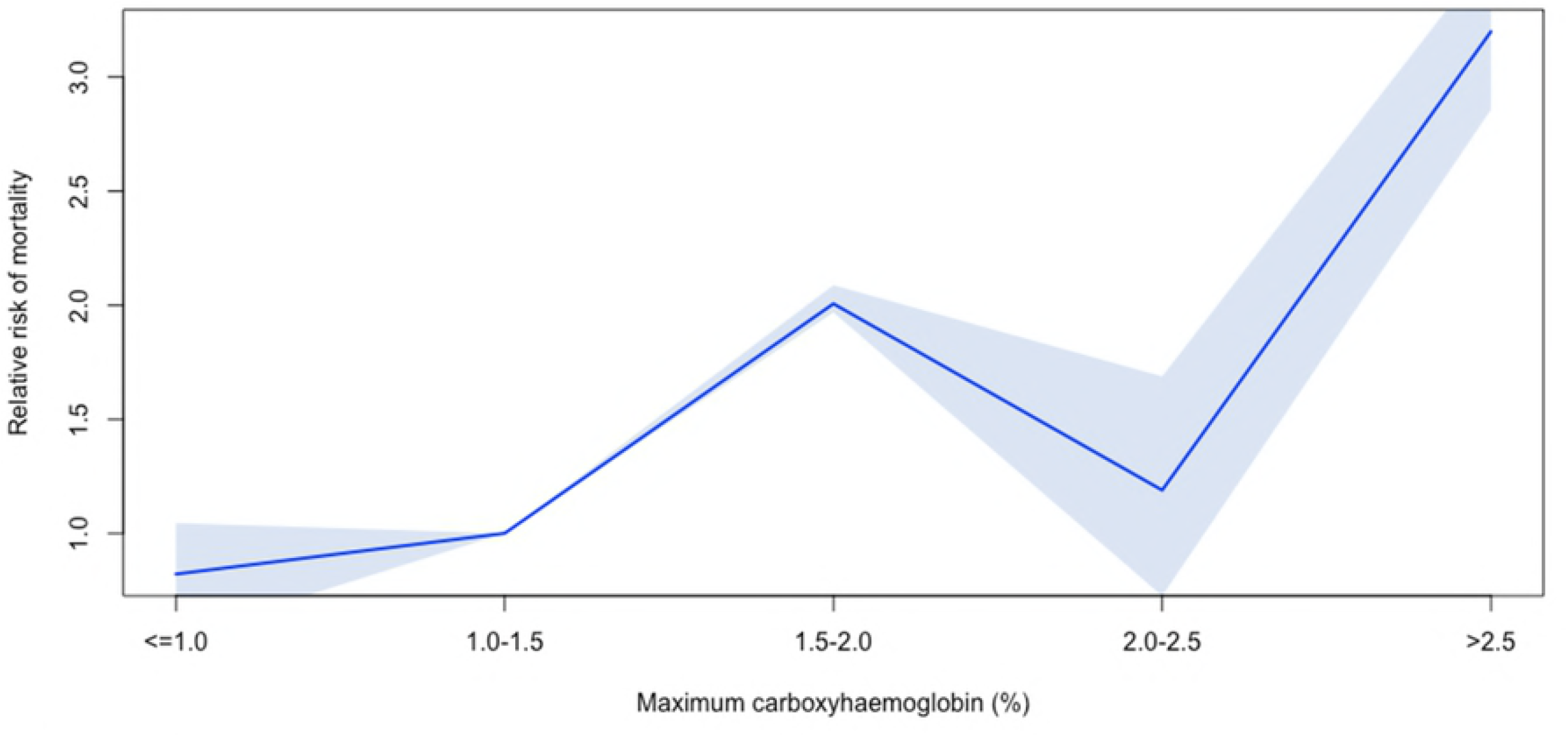
Risk adjusted mortality according to categories of maximum COHb, relative to a maximum COHb value between 1.0-1.5%. The shaded blue area represents the 95% confidence interval. Risk of mortality is adjusted for PIM-3, age and length of ventilation. The U-shaped relationship described by Melley et al in adults is not seen – COHb values <=1.0 have a non-significantly lower risk adjusted mortality relative to children with a maximum COHb between 1.0-1.5%.

Our study has several limitations. Our numbers are restricted by only including children with arterial samples, although these are children are likely to have more severe disease, and have a higher risk of mortality (U.K. baseline PICU mortality 3.8% cf 8.6% in our cohort). We excluded children who required CVVH given the risk of extra-corporeal haemolysis. However these children are likely to have greater severity of illness needing organ support, and data from them may still be of value. Our cohort did not include children post cardio-pulmonary bypass or on extracorporeal life support – this population need consideration, but are likely to have different COHb profiles and associations (similar to differences between the corresponding Fazekas and Melley cohorts). We did not consider haemoglobin levels, which had been shown to be associated with COHb levels in premature neonates [16]. However, haemoglobin levels are likely to be more variable in the early neonatal period, with greater haemolysis.

While CO has a potential protective effect through reducing reactive species mediated inflammation; we did not see evidence of this in our population. Further work may be needed to demonstrate the effect on morbidity and recovery from critical illness. If indeed CO is protective, then this may be used as a therapeutic strategy, either with the use of exogenous CO [23,24], or by the pharmacological induction of HO-1 [25].

## Acknowledgments

None

